# The BCM rule allows a spinal cord model to learn rhythmic movements

**DOI:** 10.1101/2021.11.12.467473

**Authors:** Matthias Kohler, Philipp Stratmann, Florian Röhrbein, Alois Knoll, Alin Albu-Schäffer, Henrik Jörntell

## Abstract

Animal locomotion is hypothesized to be controlled by a central pattern generator in the spinal cord. Experiments and models show that rhythm generating neurons and genetically determined network properties could sustain oscillatory output activity suitable for locomotion. However, current CPG models do not explain how a spinal cord circuitry, which has the same basic genetic plan across species, can adapt to control the different biomechanical properties and locomotion patterns existing in these species. Here we demonstrate that rhythmic and alternating movements in pendulum models can be learned by a monolayer spinal cord circuitry model using the BCM learning rule, which has been previously proposed to explain learning in the visual cortex. These results provide an alternative theory to CPG models, because rhythm generating neurons and genetically defined connectivity are not required in our model.

**Author summary:** The central pattern generator is the leading hypothesis of locomotor control in animals. There, rhythm generating neurons and genetically defined neural connectivity would form a circuit generating activity patterns suitable for locomotion. We provide a new hypothesis of locomotor control, where rhythmic patterns are learned by a Hebbian learning rule from a mechanical system that has an intrinsic tendency to oscillate.

## Introduction

Currently a central pattern generator (CPG) in the spinal cord is hypothesized to be responsible for locomotor control in animals [1]. Early experiments showed that decerebrated cats can locomote if held on a propelled treadmill [2]. Later evidence showed a reciprocal organization of inhibitory spinal interneurons [3–5], which could facilitate alternation between antagonistic muscles and between left and right limbs. Isolated spinal cords without afferents can generate locomotion patterns without rhythmic input [1, 5, 6] and such activity patterns are called fictive locomotion. All of these phenomena were ascribed to rhythm generating neurons [7, 8]. In addition, there are other neurons that belong to the locomotor circuits and aspects of the connectivity between them have been described as being dependent on their gene expression phenotype [9]. The combination of these findings have been used to generate models, which are sufficient to explain coordinated patterns of motor output thought to underlie locomotor control [9–13].

This current understanding of the spinal cord has a number of limitations. Rhythm generating mechanisms proposed for the spinal cord circuitry either rely on rhythm generating neurons or require unphysiological assumptions of properties determining network behavior in order to explain the range of observed locomotor behaviors [14]. In order to activate rhythmic activity in the spinal cord in adult animals *in vivo*, without activation of rhythmic sensory afferent feedback patterns, drug manipulation is required [15]. However without drug manipulation, spinal neurons do not intrinsically generate rhythms in adult animals *in vivo* [16]. Furthermore, the normal locomotor behavior of animals in nature is much more varied and flexible to the circumstances of the terrain than the current CPG models can explain. In most CPG models [9–13], the connectivity between neurons is assumed to be fixed, which means that the model would not be compatible with, for example, reorganized biomechanics. However, after tendon transfer surgery, cats are still capable of developing normal locomotor behavior [17]. Beyond this demonstration of flexibility of the spinal cord circuitry, the observation that “the skate spinal cord has the same interneuronal building blocks available to form the locomotor network as seen in mammals” [18] is in fact another strong indication that the spinal cord circuitry is to a large extent dependent on learning. If the genetic construction of the circuitry is the same across species that have very different biomechanical configurations, this must mean that the spinal cord circuitry is highly adaptable to the mechanics of the body it is connected to.

We propose a new model of biological locomotor control, to address the limitations of current CPG models. Our model could explain how the spinal cord acquires the capability to contribute to locomotion control. This model does not use neurons that intrinsically generate rhythm nor an initial network structure that would facilitate rhythm generation. Instead we use a generic network of neurons, without any assumptions of a priori formed connectivity, attached to a mechanical system, which has a tendency for intrinsic rhythmic movement but otherwise is unable to sustain rhythmic movement on its own because of damping. Moreover we assume that the synaptic weights of the neurons can be learned according to the BCM rule [19, 20]. This rule has been proposed to explain learning in the visual cortex and is widely considered as a good model of synaptic learning rules in the brain. The BCM rule tries to find synaptic weights so that the firing rate of the neuron has a bimodal distribution and that the average firing rate has a fixed value. In particular the BCM rule has been applied to learn from passively received input. In our model, the neurons learn from sensory input from the mechanical system, whose activation is only controlled by the output of these neurons. Hence, in our model the neurons learn from the input they themselves are responsible for creating, in a closed loop. Our model was capable of learning to control two quite different mechanical systems, one composed of two independent pendulums and one composed of one double pendulum, which are common models in control theory. A double pendulum is also the most basic model of a limb. Our results show that under these conditions and assumptions, a simplified spinal cord circuitry can adapt to different mechanical systems to generate rhythmic and alternating movement, in that respect being equivalent to current CPG models.

## Methods

### Neuron and network model

The neuron model was similar to a previous non-spiking model [21, 22]. Each neuron had an output activity, which was a time continuous voltage, called firing rate, and was modeled as follows. Let 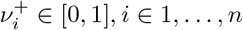 be the firing rates of neurons and external inputs connected with positive weights 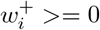 to the neurons and 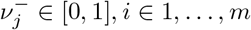 the firing rates of neurons connected with negative weights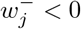. The dynamics of the potential are

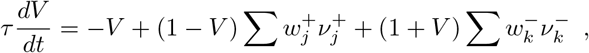

where *τ* = 5 ms. The term − *V* modeled a leak, the factors 1 −*V* and 1 + *V* modeled reversal potentials which limited the maximum and minimum voltages the neuron could have. The neurons firing rate *ν* = max(0, *V*) was computed by thresholding the voltage at zero.

The network structure was defined by the connection weights between the neurons and between inputs and neurons. Connections from a neuron to itself were not present in this model. All weights were subject to learning, as described in the next section. The weights between two distinct neurons were initialized with a random sample from 𝒰 (−0.9, 0.9) (uniform distribution). The connections between an external input and a neuron were initialized with a random sample from 𝒰 (1.5, 2.9), to somewhat bias the neurons towards external input.

### Learning

The learning model was adapted from the BCM rule [19, 20]. For each neuron with firing rate *ν* a learning threshold *ϕ* was maintained. The learning threshold is the low pass filtered version of the squared firing rate, with a time constant *τ*_*ϕ*_ = 500 ms,

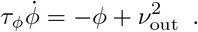

The time constant *τ*_*ϕ*_ was chosen to be substantially larger than *τ*.

For a connection with weight *w*, where a neuron with firing rate *ν*_in_ provides input to a neuron with firing rate *ν*_out_ the change of the weight was

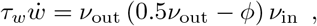

where *τ*_*w*_ = 10 s. For the BCM rule an equilibrium is obtained when the learning threshold and the voltage are both either one or zero. For the model neurons used here it is impossible to reach a voltage of one. Therefore the factor 0.5 was introduced in the above equation, which moved the equilibrium to a voltage and a learning threshold of 0.5.

### Mechanical models

We used two mechanical system models, two independent pendulums 1 A and a double pendulum 1 B. The models consisted of links connected by rotational joints, either to another link or an unmovable base. All joints had friction.

### Independent Pendulums

Let *θ*_*i*_, *i* ∈ 1, 2 be the joint angles of each of the two pendulums. In each joint a linear spring with spring constant 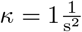 was located. The joint had friction 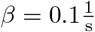. An external force *F*_*i*_ could be applied to each joint. No gravity acted on the pendulums. The dynamics of pendulum *i* was

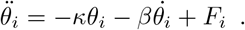

### Double Pendulum

Let *θ*_1_ and *θ*_2_ be the joint angles of the two joints in the double pendulum. Both links had length *l* = 2m, mass *m* = 1kg and the center of mass was, relative to the joint, at *l*_*c*_ = 1m. The moment of inertia of each link was *I* = 1kgm^2^ and the friction in each joint was 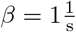. A gravity of *g* = 9.81m*/*s^2^ acted on the pendulum. The dynamics were given by

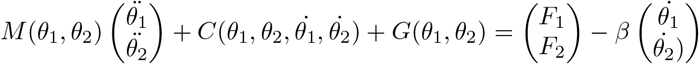

The mass matrix was

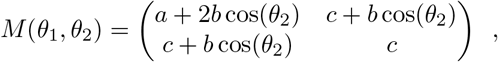

where

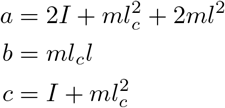

Let *h* = −*ml*_*c*_*l* sin(*θ*_2_), the Coriolis matrix was

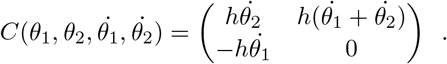

The gravity vector was

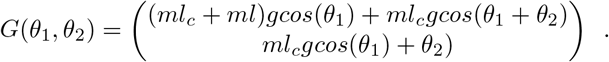

The cartesian coordinates of the endpoint of the double pendulum are

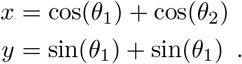

### Network and mechanics closed loop

In order for the network to control the mechanical system, the two systems were connected in a closed loop 1 C. Each neuron received as an external input the joint angles, velocities and the negative joint angles and velocities thresholded at zero and limited to one. The inputs were min(max(0, − *x*), 1) and min(max(0, *x*), 1) for all 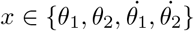. This way all inputs to the neurons are positive and negative angles and velocities are presented to the neurons on a different input than positive angles. Additional to the input from the mechanical system, each neuron received a motor command in the format of a random external input. This input was used to initiate movement in the system. It was generated for each neuron separately by random sampling from 𝒰 (0, 0.9).

The torques applied in the joints were computed from the neurons firing rates

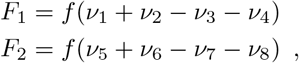

where *f* is a constant factor, which scaled firing rates of the neurons so that a torque leading to reasonable movement in the mechanical system is produced. For the independent pendulums *f* = 12 and for the double pendulum *f* = 6. If this model is to be adapted to different mechanical systems than presented here, this factor must be adapted too.

### Learning and testing

The network and mechanics were initialized so that the neuron voltages were zero and there was no movement. Weights were initialized randomly as described above. First, learning was disabled and it was tested if rhythmic movement could be generated. This was done by sampling a constant external random input (motor command) for each neuron independently, running the simulation for 100 s and then determining if rhythmic movement was generated in the second half of the simulation, as described below. The test set was composed of one hundred random combinations of motor command inputs each lasting 100 s. Then learning was enabled and the simulation was run for 2000 s. Every 1 s a new random motor command was sampled for each neuron. Finally, learning was disabled again and it was tested if rhythmic movement could be generated, using the same set of one hundred test inputs, the same way as before the learning.

In order to determine if there was rhythmic movement in a test, the joint angle time sequences were plotted. The plots were inspected if rhythmic movement was visible and if the amplitude did not decay in the second half of the test.

Additionally, to visualize the result of all tests, the autocorrelation of each joint angle sequence of the second half of each test was computed. The period length of the oscillation was taken as the position in time of the second largest local maxima of the autocorrelation. If only one local maxima was available, the period length was estimated as zero, which indicated that there was no rhythmic movement. The amplitude of the oscillation for each joint angle in each test was computed as the difference between the largest and the smallest value of the joint angle.

### Numerical methods

Each neuron, the learning and the mechanical system were simulated with an integration step size of 1 ms. The neuron dynamics were integrated with the inverse Euler method. The learning was integrated with the forward Euler method. The mechanical system was integrated with the Runge-Kutta-4-5 method. The entire simulation was implemented in C++ using the Eigen [23] and boost::odeint libraries.

## Results

We used two different pendulum systems to test if rhythmic movements could be learned using the BCM rule in neurons modeled without any intrinsic active conductances or rhythm generation (Fig 1). The pendulums were designed to hang from an attachment point, similar to a leg (Fig 1A,B). In one case, we used two independent pendulums, similar to left and right legs, which were spring-loaded with the equilibrium in the vertical orientation. The double pendulum had no springs but gravity acted on the masses of the two segments. Both pendulums had friction in the joints so that active actuation was required to generate and sustain movement. The joints had sensors that reported the joint angle and joint velocity. These sensors were provided with activation thresholds and then the sensor information was provided as input to a neural network (Fig 1C). In order to signal the joint angles and velocities in both directions, the sensor information from each joint was duplicated and half of the sensor information set was inverted before the thresholding (Fig 1C). Hence, each sensor signal was excitatory. Furthermore, each neuron received input from each of the eight sensors. The eight neurons were located in a one-layer, fully connected network, where each neuron was connected to every other neuron except for itself. The weights of all synapses were modifiable through learning. The neurons had outputs to drive movements at the joints (joint torque input), where each joint was controlled by four neurons, of which half controlled a negative torque proportional to its firing rate (in this case the thresholded voltage output of the neuron), and half controlled a positive torque. Each neuron in addition received a unique motor command input, which was a random tonic excitatory

**Fig 1.**
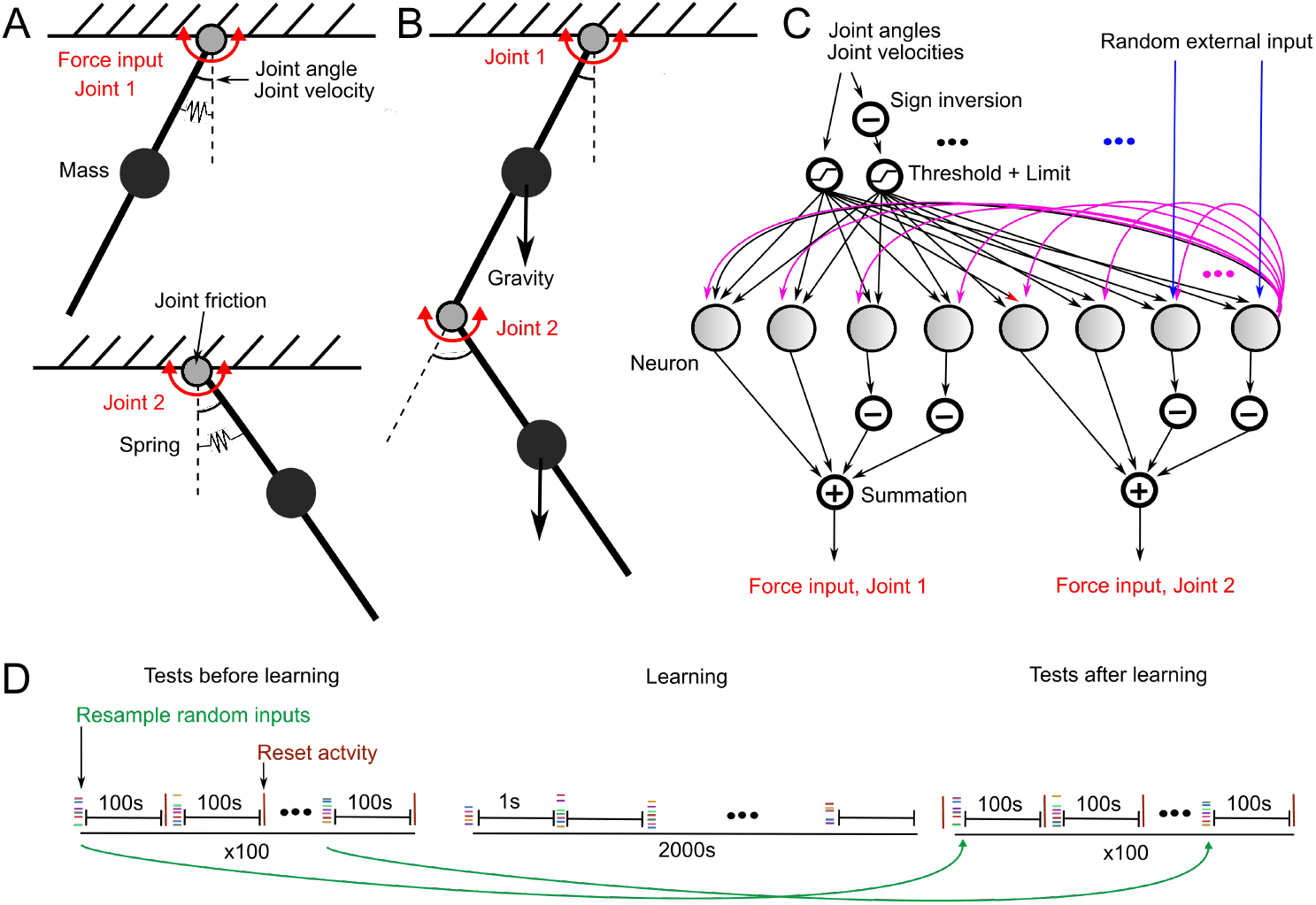
Illustration of the models. (A) Independent pendulums. (B) Double pendulum. (C) Structure of the neural network. Inputs and outputs, to and from the mechanical system (black arrows), connections between the neurons (magenta arrows) and connection from random motor command inputs to the neurons (blue arrows). Each neuron received an independent level of external input in one synapse only, whereas all inputs (sensory and internal) reached all neurons. (D) Motor commands used for network training and for testing for the presence of rhythmic movement. Each test motor command lasted for 100 s and was composed of random tonic levels of input activity in each external input. Before and after the learning, the same set of 100 tests were applied. After each test, the activity of the system, the neuron firing rates and the movements of the pendulums were reset to zero. During the learning, the random input was resampled every second and a much higher number of input combinations were applied.

### Generation of rhythmic movement

The BCM rule allowed our model to learn to control rhythmic and alternating movement in both mechanical systems (Fig 2 A-C).

**Fig 2.**
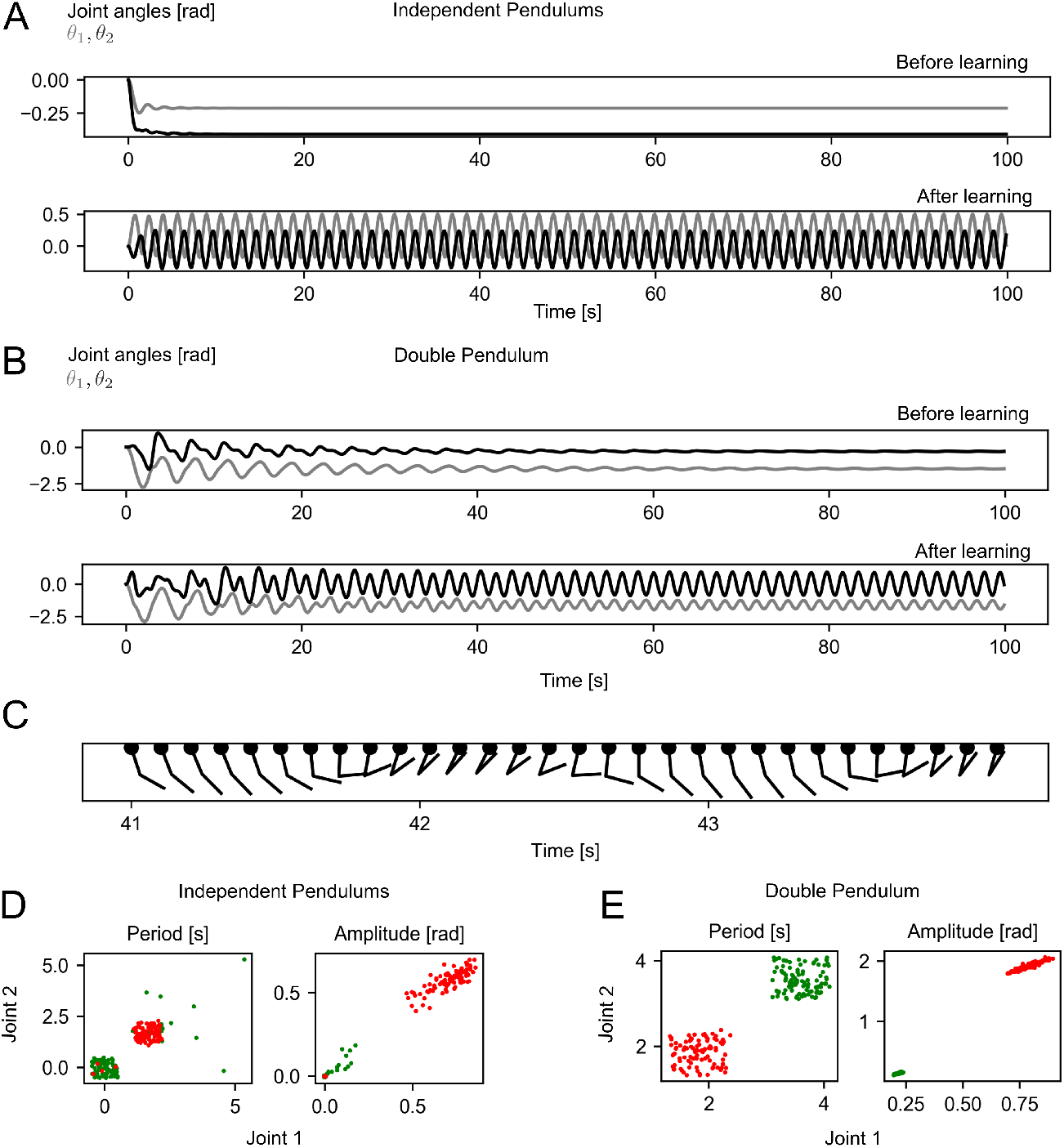
Movement patterns before and after the learning. (A, B) Examples of joint angle sequences for each mechanical system from one test before and after the learning. (C) Movement sequence of the double pendulum during one test after the learning. (D, E) Estimated oscillation period length and amplitude of each joint for each mechanical system and each test, before the learning (green points) and after the learning (red points). The points are offset in both axis by a random sample from 𝒰 (−0.5, 0.5) for the period and 𝒰 (−0.005, 0.005) for the amplitude plots in order to improve readability. input level, through a synapse whose weight was also modifiable through learning. Motor command inputs were required to elicit movement and thereby sensory feedback information. One simulation was composed of an initial test set with 100 randomized motor command input combinations, followed by a learning phase of 2000 different motor commands (Fig 1D) and finally another test set with the same motor command inputs as in the first test.

After the learning for the independent pendulums system, almost all tests showed rhythmic movement with an amplitude larger than in any test with rhythmic movement before the learning (Fig 2 D). Only few tests before the learning showed rhythmic movement at all. The distributions of the periods and amplitudes were narrow both before and after the learning, indicating that the structure of the external input (motor command) did not greatly impact the movement pattern in either case (Fig 2 D). In all tests the rhythmic movements between the two pendulums were alternating, despite no mechanical coupling between the two independent pendulums.

For the double pendulum in all tests before the learning, the system showed rhythmic movement throughout the entire test phase, but the amplitude of the movement decayed with each oscillation cycle (Fig 2B, E). The movement of the double pendulum resembled an arm repeatedly picking up and lifting an object (Fig 2 C). The trajectory, drawn out by the endpoint of the double pendulum, is shown in Figure 3C. After the learning all tests showed rhythmic movement with non-decaying amplitude. Across all tests after the learning, the movements had similar periods and amplitudes for each joint (Fig 2D), indicating rhythmic movement.

**Fig 3.**
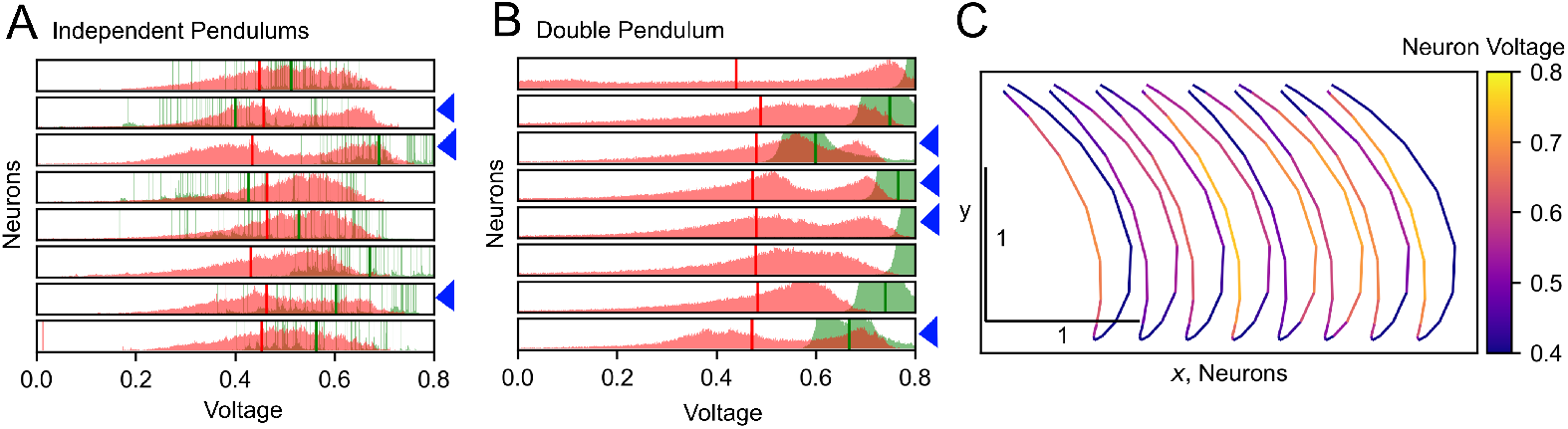
Distribution of the neuron voltages. (A,B) distributions of the neuron voltages across each neuron (one neuron per row). The voltages are accumulated over the test phases, and are shown separately for the phase before the learning (green) and after the learning (red). Vertical thick lines indicate the mean of the distributions. Note that one distribution had a mean above 0.8 before the learning. (C) The activity, or voltage, of each of the eight neurons is color coded and plotted against the position of the endpoint of the double pendulum during one oscillation cycle. Each neuron is plotted with a at an offset along the X axis.

These two experiments illustrate that the network was able to adapt to both mechanical systems and to control rhythmic movement in both of them.

### Distribution of the neuron voltages

The learning changed the distribution of the output activity levels or voltages for each neuron. In the case of the independent pendulums, voltages were distributed sparsely before the learning, because the mechanical system remained mostly at a constant position during the tests. After the learning, voltages were not sparsely distributed anymore (Fig 3 A). The grand mean of the distribution means before the learning was 0.54 with a standard deviation of 0.09. After the learning the grand mean was 0.45 with a standard deviation of 0.01, which indicates that the neurons during learning converged towards the target activity defined by the BCM rule. In the case of the double pendulum system, the distributions of the output activity of the neurons showed no sparsity before the learning because the pendulum showed rhythmic movement, but with decaying amplitude. However, the distributions had a high spread with a grand mean of 0.74 and a standard deviation of 0.07. After learning, the spread substantially narrowed down to yield a grand mean of 0.47 with a standard deviation of 0.01 (Fig 3 B).

In addition, after the learning the voltage distributions of three of the eight neurons for the independent pendulums system were clearly bimodal, in line with the expected result of the BCM learning rule. For the double pendulum system, learning instead resulted in four of the neurons having bimodal distributions (Fig 3 A,B blue arrowheads for bimodal distributions).

Figure 3C shows the activity of each neuron during one oscillation cycle of the double pendulum. Each neuron tended to be selectively active for a specific phase of the movement, but the degree of selectivity varied between the neurons. Especially the rightmost four neurons, which corresponded to the neurons controlling the second joint, were highly selectively active for one specific movement direction of the pendulums endpoint.

For both mechanical systems, the learning rule found bimodal voltage distributions for a subset of the neurons and shifted the mean firing rates of all neurons close to 0.5, which was to be expected from the BCM rule.

## Discussion

We have demonstrated that a randomly connected network can learn to generate patterns of rhythmic and alternating movement in different mechanical systems. The model of the neurons and the synaptic learning have been proposed previously [19–22]. We used them without modification, except for a shift of the equilibrium point of the learning model to adapt it to the neuron model (see below). The model was made very simple, to only demonstrate the basic principles. The pendulums served as crude approximations of animal legs. The monolayer of neurons was designed to resemble a simplified spinal cord circuitry in terms of its sensorimotor connectivity, without any separate classes of interneurons with respect to predefined sensory inputs.

Our model could potentially explain how the biological spinal cord learns to control locomotion. Note that our system for example automatically learned ‘left-right alternation’ between two mechanically independent pendulums (Fig 2C). Our model could potentially explain how the biological spinal cord controls locomotion. To map our model to an envisaged ontogenetic development of the spinal cord circuitry in an animal, initially neurons would need to form connections with each other at random and they would need to receive random connections from the sensory afferents. Also the connectivity from the spinal interneurons to the motoneurons, each innervating a specific muscle, would be random. The activity of the spinal neurons would stimulate force generation by the muscles, which through the biomechanical configuration of the body would trigger subsets of preferred movements. In a system that is inclined to rhythmic movement, like the systems used here, this would lead to some rhythmic movement and the sensory input to the spinal neurons, that results from that movement, would also be rhythmic. From this the neurons can learn, so that they have stable firing rates and can provide input to other neurons or to the muscles. This would lead to the muscles having rhythmic activation, which which would result in rhythmic movements as shown here. Inhibitory connections could allow neurons to reduce the activity of other neurons, allowing alternating activity between neurons, so that movement patterns where antagonistic muscles are active alternatingly could form.

Moreover inhibitory and excitatory connections between neurons could allow the distribution of neuron activity to change during a movement, where the distribution of muscle activity evolves during the movement. Both alternating activity and shifts in neuron activity could be facilitated by a learning rule that finds synaptic weights so that each neuron is selectively active for different phases of a movement. In reality the brain would have to select motor command inputs to the spinal cord that result in more varied movements that would be suitable to fulfill the goals of the brain. Such motor command inputs, in turn, could influence the movement patterns from which the spinal cord circuitry structure could learn.

Our model could also help to find and control oscillatory modes of general multi-body mechanical systems. Intuitively, modes are intrinsic movements of mechanical systems, which are determined by the ability of a system to store and release energy [24]. For the double pendulum system, our model controlled a movement similar to a so called energy invariant mode, where the pattern of movement is the same regardless of how fast the movement is [25]. The double pendulum supports a greater variety of modes and developing controllers for them would enable robots to achieve energy efficient movement [24]. Future work could be to find neural networks and learning rules able to learn to control these modes.

Compared to CPG models of locomotion, our model did not use rhythm generating neurons and the initial connectivity of the network was random. It has been proposed that central pattern generators are composed of rhythm generating neurons [1] that are connected in a network determined by the genetic expression of the neurons [13]. Our model assumes that learning exists in the spinal cord, similar to what has been proposed for the brain. While previous models of locomotion without rhythm generating neurons have been proposed, they either had manually adjusted synaptic weights [26] or an immutable network structure facilitating locomotion [27]. Overall, our results demonstrate in principle how neural network behavior, which could explain phenomena compatible with the observations underlying the notion of central pattern generators, can arise through learning without any genetically programmed network connectivity or rhythm generating neurons.

However our model has a few limitations that current CPG models do not have. First, the amount of experimental publications on learning in the spinal cord is less than the amount of experimental publications on CPGs. Also, our model did not address the fictive locomotion phenomenon.

During the learning phase, a more organized external input than the independent random signals that we used could lead to a greater variety of random movement from which the network structure could be learned. In a developing animal, such semi-organized but still predominantly random movements could be generated by the spontaneous internal activity of the brain [28, 29], spontaneous activity of spinal neurons and by random muscle twitching.

If the model is able to learn rhythmic movement or not is critically dependent on the so called force factor. This factor is the gain, that converts the firing rates of the neurons into a force in the mechanical systems. Different mechanical systems, depending on their masses, frictions and springs, require different forces to move. If the factor is too small, only little movement is generated and it is impossible to learn rhythmic movement. If the factor is too large, only erratic movement is generated, also making it impossible to learn. For the double pendulum, it could be of advantage to use different force factors for each joint, because the total masses that move around each joint are different. In biology muscles and spinal cord must develop during the same time. It could be that muscles must adapt to the inputs from the spinal cord.

The models of the mechanical systems were chosen to be similar to the ones used in control theory and robotics as models of robotic and biological limbs for the sake of simplicity. Models of actual biomechanical systems such as cat legs or entire lamprey bodies would be an important test that was not considered here.

The BCM rule has been proposed to explain how neurons in the brain could learn to be selective for features in passively received visual sensory inputs. Here the BCM rule works in a model where input is actively influenced by the model itself. The key features of the rule, that make this possible are stabilization of the firing rates of the neurons and optimization for a bimodal firing rate distribution. A stable firing rate in our case helped to provide an, on average, limited force input to the mechanical systems. Bimodal firing rate distributions can help make neurons selective for certain phases of a movement and provide activity during this phase.

Moreover, we combined the BCM rule with a nonlinear neuron model, where the maximum firing rate is always smaller than one. If we had used the BCM rule as given in the references [19, 20, 30], we would have weights that converge to infinity, because this formulation has an equilibrium point when the firing rate is one. In one study [30] weight growth was counteracted by a learning rate depending on the learning threshold. Instead we shifted the equilibrium point to a firing rate that can be attained by the model neurons.

Clearly limitations in this model and in existing CPG models must be addressed to find an accurate model of the contribution of the spinal cord to rhythmic movement generation. In biology, it is likely that the mechanisms provided in previous CPG models, in combination with developmental, adaptive, and learning processes such as the ones described here, are contributing to an integrated whole.

## Acknowledgments

This research received funding to AK from the German Aerospace Center under the research and development contracts D / 572 / 67 24 68 70, D / 572 / 67 27 00 80, the European Union’s Horizon 2020 Framework Programme for Research and Innovation under the Specific Grant Agreements No. 720270 (Human Brain Project SGA1), 785907 (Human Brain Project SGA2), 945539 (Human Brain Project SGA3).

Employees of the German Aerospace Center are coauthors of this study.

